# TPX2 Amplification-Driven Aberrant Mitosis in Long-Term Cultured Human Embryonic Stem Cells

**DOI:** 10.1101/2021.02.22.432205

**Authors:** Ho-Chang Jeong, Young-Hyun Go, Joong-Gon Shin, Yun-Jeong Kim, Min-Guk Cho, Dasom Gwon, Hyun Sub Cheong, Haeseung Lee, Jae-Ho Lee, Chang-Young Jang, Hyoung Doo Shin, Hyuk-Jin Cha

## Abstract

Although human embryonic stem cells (hESCs) are equipped with highly effective machinery for the maintenance of genome integrity, the frequency of genetic aberrations during long-term *in vitro* hESC culture has been a serious issue that raises concerns over their safety in future clinical applications. By passaging hESCs over a broad range of timepoints, we found that mitotic aberrations, such as the delay of mitosis, multipolar centrosomes, and chromosome mis-segregation, were increased in the late-passaged hESCs (LP-hESCs) in parallel with polyploidy compared to early-passaged hESCs (EP-hESCs). Through high-resolution genome-wide approaches and by following transcriptome analysis, we found that LP-hESCs with a minimal amplicon in chromosome 20q11.21 highly expressed *TPX2* (targeting protein for Xklp2), a key protein for governing spindle assembly and cancer malignancy. Consistent with these findings, the inducible expression of TPX2 in EP-hESCs reproduced aberrant mitotic events, such as the delay of mitotic progression, spindle stability, misaligned chromosomes, and polyploidy. This data suggests that the amplification and increased transcription of the *TPX2* gene at 20q11.21 could contribute to an increase in aberrant mitosis due to altered spindle dynamics.

## Introduction

While the machinery of genome integrity maintenance is well-developed in human embryonic stem cells (hESCs) (1), genetic abnormalities ranging from full chromosome aneuploidy to single point mutations are relatively common in human pluripotent stem cells (hPSCs) when they are propagated in *in vitro* culture (2, 3). Hence, genetic aberration in hPSCs during *in vitro* maintenance has remained a serious hurdle for future clinical applications (4) because of the uncertainty of the biological consequences of genetic aberrations in hPSCs (5). Indeed, unexpected genetic mutations identified in induced pluripotent stem cells (iPSCs) halted the second human clinical trial of iPSC-based cell therapy (6).

Through massive array comparative genomic hybridization (aCGH), the sub-chromosomal amplification of the 20q11.21 locus has been reported to be the most frequent copy number variant (CNV) with normal a karyotype (4,7–9) derived from *in vitro* culture because the gain of 20q11.21 does not occur in normal embryos (10). In particular, *BCL2L1* in 20q11.21 has been identified as a driver mutation for survival advantages in cultures (11, 12), which also results from an independent loss-of-function mutation in p53 (13). Other than the survival advantage conferred by the amplification of *BCL2L1* or the *TP53* mutation, the other genetic alterations have not been closely characterized, with a few notable exceptions (14), although various abnormal behaviors, such as tumor formation (15, 16) and impaired differentiation (17), in their progenies or hPSCs themselves were reported in animal models. Although CNV, including 20q11.21, occurs even in early passage (14, 18), the incidence of abnormal karyotypes may occur in a prolonged culture (19), suggesting that the induction of a set of gene(s) by CNV would lead to further chromosomal aberrations (20).

Trisomy of chromosomes 12 or 17, which is frequent in embryonal carcinomas (21), is commonly found in hPSCs due to the survival benefit of these abnormalities in culture (20). Recent studies demonstrated that aneuploid hPSCs (with trisomy in chromosome 12) showed increased proliferation and impaired differentiation *in vivo* (12, 17) and also developed tumors when differentiated cells were transplanted (15,16,22). However, to date, further molecular mechanisms underlying chromosomal instability (CIN) upon *in vitro* propagation have remained largely unknown, with the exception of a few studies that have revealed that reduced serum response factor (SRF) expression (23) or escape from mitotic cell death upon mitotic errors is responsible for ongoing CIN in aneuploid hPSCs (24).

A targeting protein for Xklp2 (TPX2) located in 20q11.21 has been studied with respect to mitosis-regulating microtubule nucleation, spindle assembly, and centrosome separation during mitosis. It has also been shown to play a role in cancer (25) by activating Aurora A kinase (26). The genetic abrogation of *TPX2* in mouse embryos leads to early embryonic lethality due to the incorrect formation of bipolar spindles and the mis-segregation of chromosomes (27), whereas the ectopic expression of *TPX2* is sufficient to induce polyploidy in a cancer cell model (28). Considering the strong association of TPX2 expression with CIN in cancer (29), and with poor patient survival in various forms of cancers (25), the deregulation of TPX2 is a putative driver of CIN by accelerating aberrant mitotic spindle dynamics and improper chromosome segregation.

Here, by late-passage hESCs (LP-hESCs), which showed aberrant mitotic events such as multipolar spindle, chromosome mis-segregation, and polyploidy compared to early passage hESCs (EP-hESCs), we identified TPX2, one of the genes in 20q11.21, as a putative driver for aneuploidy in hESCs, likely due to its roles in the stabilization of mitotic spindles and consequent aberrant mitosis.

## Materials and Methods

### Cell culture

hESCs (H9; Wicell Research Institute) were maintained in mTeSR1 medium (StemCell Technologies) on plates coated with Matrigel (Corning, #354277) diluted at 1:80 in hESC basal medium (DMEM/F12 supplemented with 1% non-essential amino acids, 0.1% β-mercaptoethanol, and 0.1% gentamycin, Gibco) for feeder-free conditions. The medium was replaced every day up to passaging, and the cells were enzymatically dissociated using a dispase solution (Gibco, #17105041).

### Construction of plasmid and generation of stable hESC lines

To construct the TPX2-WT or rtTA expressing piggybac plasmid, EGFP-tagged human TPX2-WT of pcDNA3-EGFPC-hTPX2-WT vector, respectively, was cloned into piggybac plasmid pB-TET. Two piggybac vectors were transfected into H9 EP-hESCs by electrophoration (NEPA), and selected using a G418 (Sigma-Aldrich, #A1720) 3 days later. After the selection, EGFP-tagged TPX2 expressing hESCs with minimal dose of doxycycline (Sigma-Aldrich, #D9891) were selected using a cell sorter (BD FACSAria II), and single cell cloning was performed.

### RNA extraction, quantitative real-time PCR, and genome-wide gene expression profiling

Real-time PCR was performed using TB green Premix Taq (Takara, #RR820A) on a LightCycler 480 Instrument II (Roche). Primers are listed in the Supplemental Materials. Total RNA was isolated using Easy-blue reagent (Intron, #17061) according to the manufacturer’s instructions. Real-time PCR was performed using TB green Premix Taq on a LightCycler 480 Instrument II (Roche). Primers are listed in the Supplemental Materials. To construct a sequencing library, we used a TruSeq Stranded mRNA Library Prep Kit (Illumina, San Diego, CA). Briefly, the steps for the strand-specific protocol are as follows: first strand cDNA synthesis; second strand synthesis using dUTPs instead of dTTPs; end repair, A-tailing, and adaptor ligation; and PCR amplification. Each library was then diluted to 8 pM for 76 cycles of paired-read sequencing (2 × 75bp) on an Illumina NextSeq 500 per the manufacturer’s recommended protocol. Sequence reads were aligned to the reference genome (UCSC hg19), and the mapped counts per gene were quantified using STAR. The raw counts were normalized to counts per million (CPM) based on the trimmed mean of M-values (TMM) normalization method using the R package “edgeR.” Quality check and subsequent trimming (removal of adapters and low-quality bases) of raw RNA-seq fastq files were conducted via FastQC and TrimGalore, respectively (https://www.bioinformatics.babraham.ac.uk/). The reads were mapped to the GRCh38 reference genome using the STAR aligner (v2.7.3a). Transcripts per million (TPM) and expected read counts were calculated using RSEM v.1.3.3. with assembly GRCh38.84. Assessment of differences in gene expression between groups was conducted using the R package “DESeq2.” DEGs were selected with |log_2_fold-change|>1 and a false discovery rate (FDR) of < 0.01. To identify genomic loci containing the most DEGs, an over-representation analysis was performed using a hypergeometric test with the positional gene sets from the Molecular Signatures Database (MSigDB, https://www.gsea-msigdb.org/gsea/msigdb/). After FDR-based multiple testing correction, an enrichment score for each genomic locus was defined as -log_10_ Q-value. For GO analysis, GSEA was performed using the GO BP gene sets from MSigDB via the R package “fgsea.” Significantly upregulated or downregulated GO terms were selected with an adjusted *P*-value of < 0.01 and a |normalized enrichment score (NES)| of > 2.

### Determination of copy number variations

Whole-genome genotyping was performed using the Illumina HumanOmni1-Quad Beadchip (Illumina) containing 1,140,419 genetic markers across the human genome. Samples were processed according to the specifications of the Illumina Infinium HD super assay. Briefly, each sample was whole genome amplified, fragmented, precipitated, and re-suspended in an appropriate hybridization buffer. Denatured samples were hybridized on a prepared BeadChip for a minimum of 16 h at 48°C. Following hybridization, the bead chips were processed for the single-base extension reaction, stained, and imaged on an Illumina iScan system. Normalized bead intensity data for each sample were loaded into the GenomeStudio software package (Illumina). Ratios of signal intensity were calculated using the Log R Ratio (LRR: logged ratio of observed probe intensity to expected intensity; any deviations from zero in this metric are evidence for copy number change) and allelic intensity was determined by the B allele frequency for all samples. Values were exported using Illumina GenomeStudio. Analysis for structural variants was performed using the sliding window approach (window size 10).

### Whole-exome sequencing

Whole-exome sequencing was performed using the Ion Proton Platform (Life Technologies). Briefly, 100 ng of gDNA was used for AmpliSeq exome amplification, according to the manufacturer’s protocol. The final sequencing libraries were inspected and quantified using the Bioanalyzer 2100 Instrument and DNA HS kit (Agilent Technologies). All libraries were diluted to 100 pM working solutions and then pooled as needed to perform the template preparation on an Ion OneTouch 2 according to the manufacturer’s protocols. Template was prepared using the Ion PI^TM^ Template OT2-200-Kitv2 on the Ion OneTouch^TM^ 2 System. Templated Ion Sphere Particles were then enriched for positive ISP using the Ion OneTouch ES, and sequencing was performed using the Ion PI^TM^ Chip-Kit v2 and Ion PI^TM^ Sequencing200-Kitv2 on the Ion Proton^TM^ Sequencer. Sequencing data were processed using the Torrent Suite^TM^ Software (ver.4.0.2) on a Torrent Server. Sequences were aligned against the reference genome (GRCh37/hg19) using the Genomebrowser (DNA Nexus).

### Spindle dynamics assay

To measure dynamics of spindle, microtubule depolymerization assay and microtubule repolymerization assay were performed as previously described (30). For depolymerization assay, cells were incubated with 1µg/mL of nocodazole and fixed with ice-cold methanol (MtOH) time dependent-manner before immunofluorescent. For repolymerization assay, cells were incubated with 1µg/mL of nocodazole, and wash-off and fix with ice-cold MtOH time dependent-manner before immunofluorescent. Mitotic spindle fluorescence intensity of metaphase cells, determined by IFC with anti-β-tubulin antibody (DSHB, #Q-E7-S) was quantified.

### Flow-cytometry

For antibody stained flow cytometry, cells were washed twice with PBS and were fixed with fix-permeabilization solution (BD bioscience, #554722). Cells were washed with permeabilization-wash solution twice (BD bioscience, #554723) and stained with primary antibodies in 3% BSA solution 1hour. 2^nd^ antibodies were used for conjugated 1hour. For measuring DNA contents, cells were fixed with ice-cold ethanol (EtOH), and washed with PBS twice. 1ug/mL of RNase A were treated in 37C 30min, and 50µg/mL of propidium iodide (PI) (Sigma-Aldrich, #P4170), 7-AAD (BD bioscience, #559925), or 2µg/mL of Hoechst33342 was stained 30min at dark.

### Immunoblotting and immunofluorescent

Immunoblotting and immunofluorescence assay were performed as described previously. Antibodies for TPX2 (#8559), phospho-Aurora-A (#3079), and Lin28A (#8706) were purchased from Cell Signaling Technology. Antibody for and β-tubulin (#Q-E7-S), was purchased from Developmental Studies Hybridoma Bank (DSHB). Antibodies for GAPDH (#SC-47724), α-tubulin (#SC-23948), were purchased from Santa Cruz biotechnology. Antibody for phospho-Histone H3 (#14955) was purchased from Abcam. Nucleus staining reagent 4’,6-diamidini-2-phenylindole (DAPI, #D1306) and Hoechst33342 (#H21492) were purchased from Thermo Fisher Scienfitic.

### Statistical analysis

Graphical data are presented as mean ± standard deviation (SD). Statistical significance for more than three groups was determined using one-way or two-way analysis of variance (ANOVA) following a Tukey multiple comparison post-test. Statistical significance between the two groups was analyzed using unpaired Student’s t-tests. Statistical analysis was performed with GraphPad Prism 8 software (https://www.graphpad.com/scientific-software/prism/). The statistical significance was assumed to be ns: not significant, \**p* < 0.05, \*\**p* < 0.01, and \*\*\**p* < 0.001.

## Results

### Aberrant mitosis and polyploidy in long-term cultured hESCs

To investigate whether long-term propagation affects the chromosomal integrity of cultured hESCs, we took advantage of four variants of H9 hESC lines with various passage numbers (P1: 40s, P2: 100s, P3: 200s, and P4: 300s). Of note, these clones showed identical short tandem repeat profiles compared to the control H9 hESCs (Fig. S1A), implying that they were derived from the same embryonic genetic background. It is noteworthy that a clear survival advantage resulting from culture adaptation occurs in hESCs over 200 passages (P3 and P4 hESCs: LP-hESCs) (12). Considering a recent study that mitotic stress survival (31) is closely associated with gaining aneuploidy (24), we first monitored mitotic progression. Interestingly, hESCs over 200 passages (P3 and P4 hESCs: LP-hESCs) showed aberrant chromosome segregation during mitosis (Fig. 1A), high incidences of chromatin bridges (Fig. 1B), and cytokinesis failure (Fig. 1C). Aberrant mitosis in LP-hESCs can be observed in real-time images (Movie S1A-D). Notably, mitotic progression was significantly retarded in P4 hESCs (Fig. 1D). Consistent with previous studies that excessive numbers of centrosomes were closely linked to improper chromosome segregation (32), supernumerary centrosomes were more frequently observed in P4 hESCs compared to P1 hESCs (Fig. 1E).

**Figure 1.**
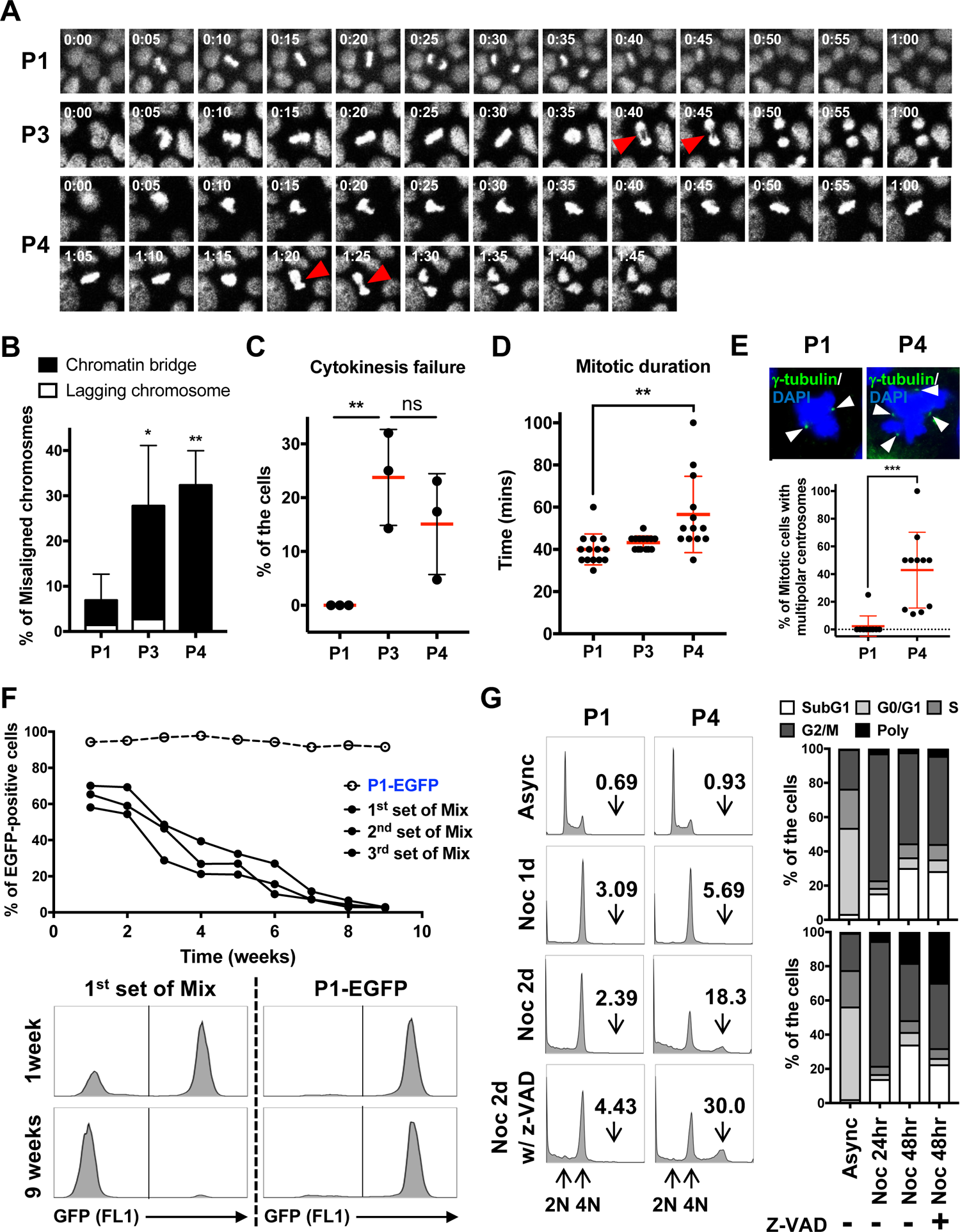
Aberrant mitosis and polyploidy in long-term cultured hESCs. (A) Time-lapse images for mitotic cells in LP-hESCs, Images were captured every 5 min and arrowhead indicates chromatin bridge. (B-C) Percentages of mitotic cells showing misaligned chromosomes (B) or cytokinesis failure (C) were quantified. (mean ± SD; n = 3, not significant: ns, \**p* < 0.05, and \*\**p* < 0.01) (D) Mitotic duration from nuclear envelop breakdown to telophase was determined. (mean ± SD; n = 14, \*\**p* < 0.01) (E) Immunofluorescent images of multipolar centrosomes in P4 hESCs, scale bar = 10µm (upper panel), Cells were stained with γ-tubulin (green) to visualize centrosomes (arrowhead) and counterstained with DAPI for chromosomes. Percentages of mitotic cells showing multi-centrosomes were represented as a scatter plot (bottom panel, mean ± SD; n = 10, \*\*\**p* < 0.001). (F) Growth competition analysis of P4 hESCs with P1 hESCs expressing EGFP, Cells were differently mixed (three independent set of mixture) as shown in 1 week, and then cultured for additional 8 passages. EGFP-positive cells were measured by flow cytometry (bottom panel) and graphically presented in upper panel. (G) Histograms of DNA contents analyzed by flow cytometry, Cells were treated with nocodazole (100ng/ml) for indicated time points in the absence or presence of z-VAD-FMK (20µM), and then the percentages of polyploid populations (>4N) were represented in left panel. Cell cycle distribution at the different phases was shown in right panel.

One clear biological effect observed in hESCs after long-term *in vitro* culture is the growth advantage (11) marked by increased self-renewal, proliferation, and resistance to apoptosis, which are referred to as “features of culture adaptation” (20). Indeed, our established P3 and P4 hESCs were progressively dominant in cultures when mixed with GFP-expressing P1 hESCs (Fig. 1F and S1B). This growth advantage might result from the acquisition of robust resistance to apoptosis (Fig. S1C) rather than increased self-renewal (Fig. S1D) or proliferation (Fig. S1E). As shown in Figure 1G, the polyploid population identified after mitotic arrest using nocodazole (Noc) was significantly enriched in P4 hESCs. In particular, the clear induction of polyploidy was observed in P4 hESCs when mitotic cell death was blocked with z-VAD-FMK, a pan-caspase inhibitor. This data clearly supports the idea that survival from the mitotic stress allows aneuploidy as previously described (24). Thus, the survival advantage acquired by “culture adaptation” in hPSCs, which leads not only to resistance to cell death but also to escape from mitotic death, contributes to the increase of polyploidy in LP-hESCs.

### Recurrent gain of the 20q11.21 genomic locus in long-term cultured hESCs

Given that 25% of hESC lines with normal karyotypes contain altered CNVs, which are defined as gains or losses of a relatively small region of a genome (9), our established cell models would exhibit this aberrant CNV profile, conferring the distinct cellular characteristics shown in Figures 1 and S1. To this end, a high-resolution single nucleotide polymorphism (SNP) array was performed in four variants of the H9 hESC lines, and the genome profile of P1 hESCs was used as a reference genome. As shown in Figure 2A, numerous gains or losses of CNVs were clearly detected in P2, P3, and P4 hESCs (i.e., 5 gains and 19 losses in P2, 6 gains and 4 losses in P3, and 14 gains and 7 losses in P4). The detected CNVs ranged in size from 479 bases to 3.10 Mb, and the average size of CNVs in P2, P3, and P4 hESCs was 55.28, 585.99, and 476.29 kb, respectively. More importantly, large CNVs (over 100 kb) were exclusively observed in P3 and P4 hESCs (Fig. 2B).

**Figure 2.**
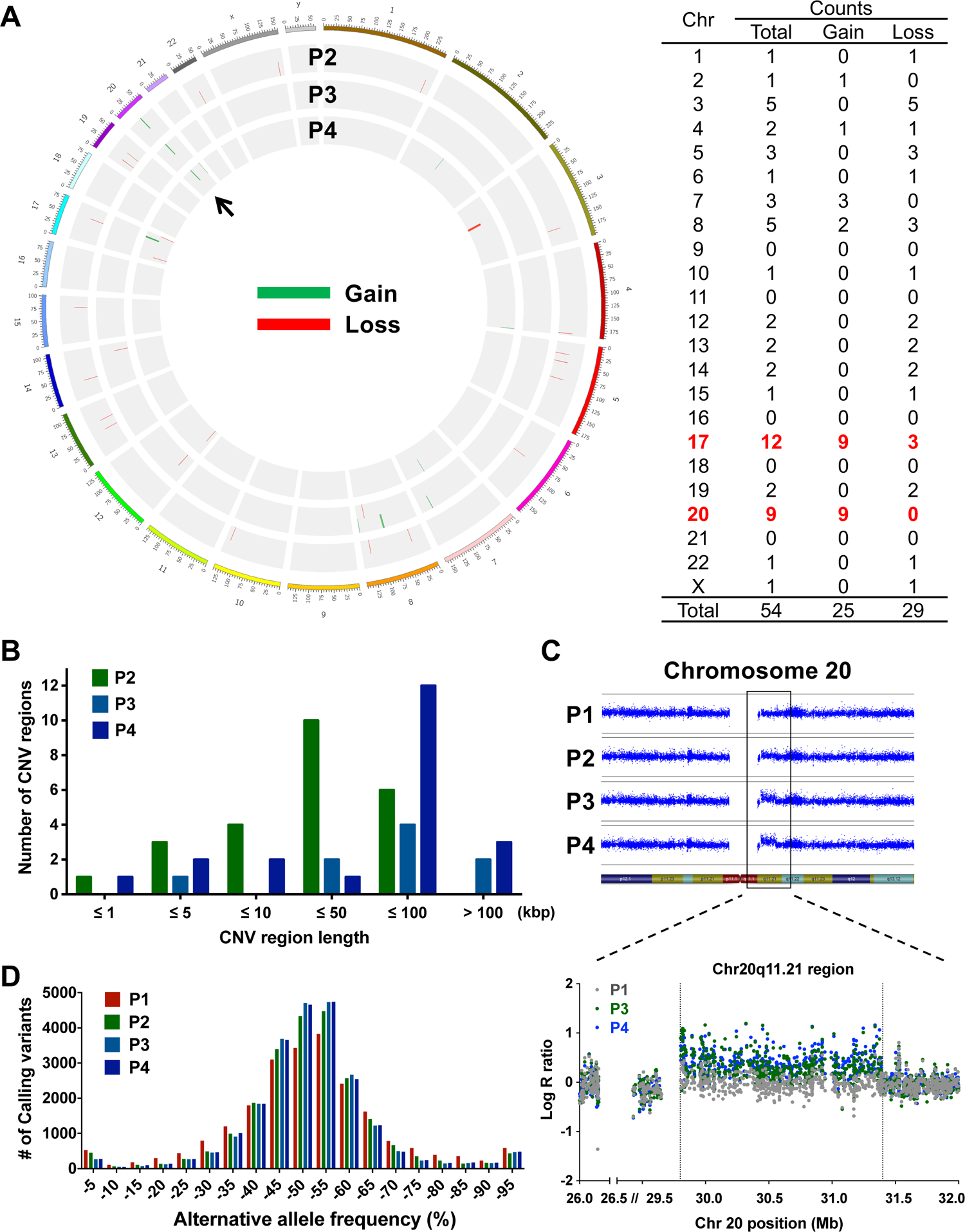
Recurrent gain of the 20q11.21 genomic locus in long-term cultured hESCs. (A) CIRCOS plot for gain (green) or loss (red) of CNVs detected by SNP array (left panel), Human 22 and XY chromosomes were represented as an outer track with each tick indicating 5 Mbps. Each circle from the outer to inner track represents the copy-number profiles of the P2, P3, and P4 hESCs. Genomic profiles of P1 hESCs were used as a reference genome and arrow indicates a commonly amplified locus. Total number of CNVs at each chromosome was represented in right panel and the chromosomes having the most CNVs were marked in red. (B) Frequency of CNVs was determined according to CNV region length. (C) Log R ratio (LRR) plots for chromosome 20 (upper panel), The intensity of each genetic probe was represented in blue-filled circle. The LRR plots for a 20q11.21 locus (about 1.5 Mbps sub-chromosomal region) were magnified in bottom panel. (D) Frequency of allelic fraction values (alternate to total read count) for 34,118 point-mutations determined by whole-exome sequencing.

The recurrent chromosome abnormalities that most commonly result from *in vitro* culture adaptation of hPSCs have been identified as the gain of chromosomes 1, 12, 17, 20, or X (20). Therefore, the repeated CNVs (i.e., gains or losses) in P2, P3, and P4 hESCs would be predicted to exist among the above chromosomes. To our surprise, the 20q11.21 genomic locus was commonly amplified in P2, P3, and P4 hESCs (Fig. 2C), and the gain of the 17q24 genomic locus was also detected in P4 hESCs (Fig. S2A and B). It is noteworthy that the gain of various loci of 17q has also frequently been associated with chromosome abnormalities (9).

Of note, given that point mutations are mostly observed in alternative allele frequencies (AAFs) at a rate of around 50% (33), the high calling variants in P2, P3, and P4 hESCs found in AAFs ranging from 40 to 60% indicates that their chromosomes are genetically unstable compared to those of P1 hESCs (Fig. 2D). Taken together, these findings indicate that long-term cultured hESCs appear to undergo continued genetic alterations, in which the additional gain of 20q11.21 allows the clones to survive against selective pressures on genomes.

### Perturbed microtubule dynamics in long-term cultured hESCs

To explore the influence of recurrent CNVs on gene expression, RNAseq was performed on the four hESC lines (P1–P4). We observed distinct overall gene expression patterns between the EP (P1 and P2)- and LP (P3 and P4)-hESCs (Figs. 3A and S3A). Differentially expressed genes (DEGs) also increased dramatically when passage number exceeded 200 compared to EP-hESCs (Figs. 3B and S3B). To focus on dysregulated genes in LP-hESCs, DEGs were selected by comparing LP-hESCs and EP-hESCs (Table S1). As expected, based on the observed gain of 20q11.21 in LP-hESCs, the top enriched genomic position of upregulated DEGs was 20q11.21, followed by Xq22 and Xp11, while no significant enrichment was detected in downregulated DEGs (Fig. 3C and Table S1). The expression of the 18 upregulated DEGs located on 20q.11.21 increased gradually over the passages (Fig. 3D), supporting a tight relationship between 20q11.21 CNV and the corresponding gene expression.

**Figure 3.**
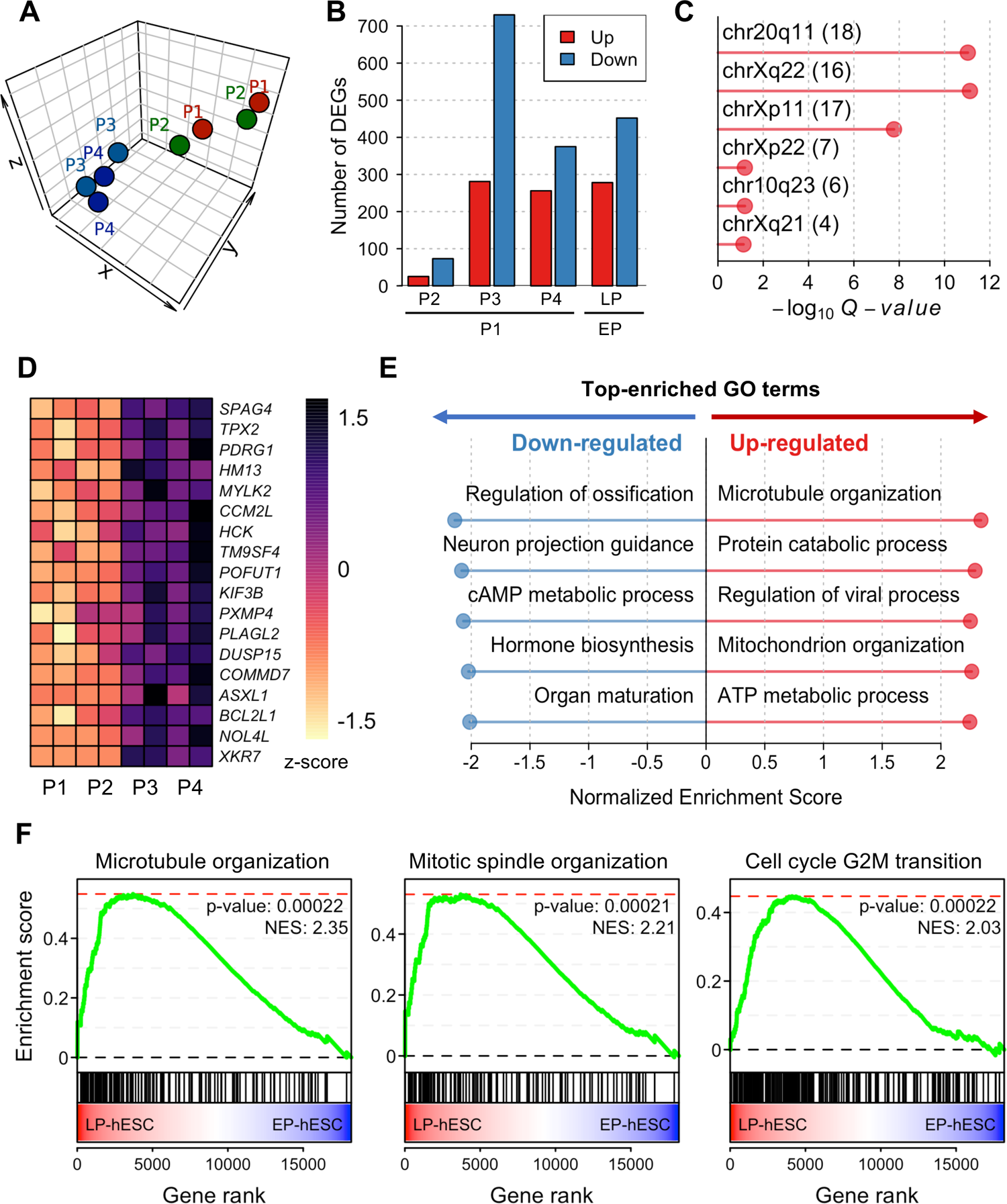
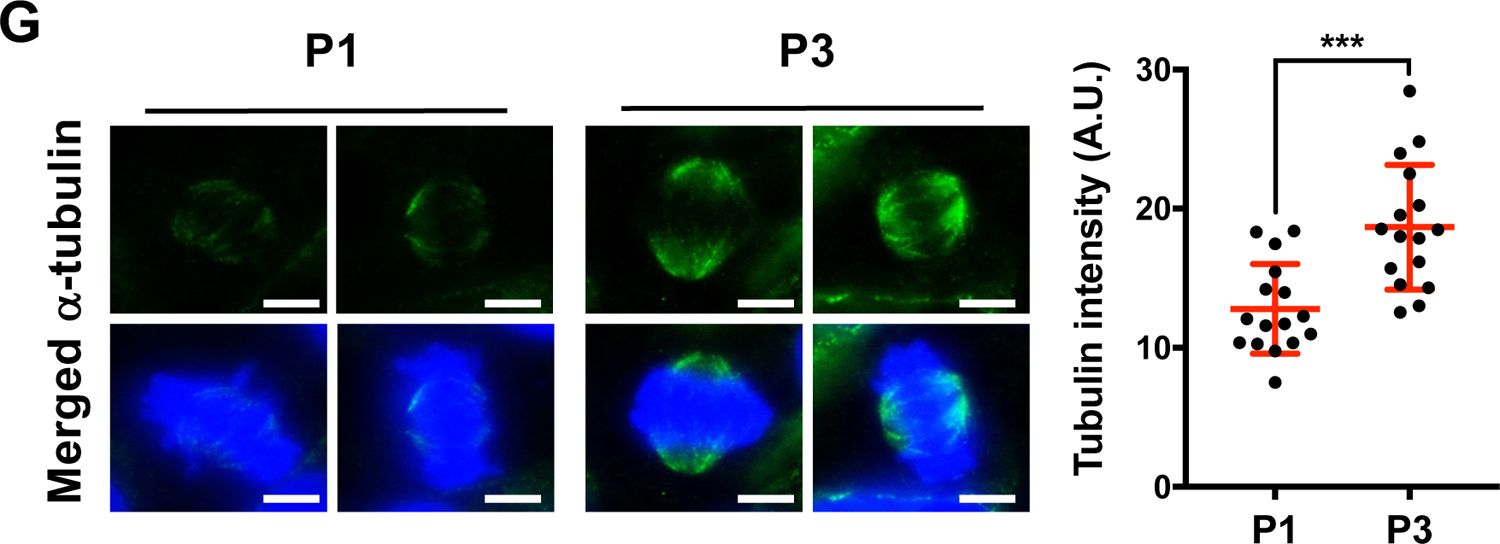
Perturbed microtubule dynamics in long-term cultured hESCs. (A) T-Distributed Stochastic Neighbor Embedding (t-SNE) clustering based on gene expression profiles of four hESC lines (P1-P4) in duplicate. (B) The number of DEGs in P2, P3, and P4 hESCs compared to P1 hESCs and LP-hESCs compared to EP-hESCs. (C) The top enriched genomic loci for upregulated DEGs in LP-hESCs. (D) Expression patterns of the 18 genes located on chromosome 20q11.21. (E) Significantly upregulated or downregulated GO terms in LP-hESCs compared to EP-hESCs. Representative GO terms were selected from Fig. S3C. (F) GSEA plots showing the enrichment of GO terms (GO:1902850, GO:0007052, GO:0044839) in a ranked list of genes differentially expressed between LP-hESCs and EP-hESCs. (G) For immunofluorescent images of microtubules (scale bar = 10µm), cells were stained with α-tubulin (green) and counterstained with DAPI for chromosomes. The fluorescent intensity was measured and represented as a scatter plot (mean ± SD; n = 16, \*\*\**p* < 0.001).

To examine functional dysregulation in LP-hESCs, gene set enrichment analysis (GSEA) (34) with gene ontology (GO) was carried out. The most upregulated functions were classified into five categories: microtubule organization, protein catabolic processes, regulation of viral processes, mitochondrion organization, and ATP metabolic process (Figs. 3E and S3C and Table S2). Interestingly, *KIF3B* and *TPX2* in 20q11.21 contributed most to the enrichment signals for microtubule-, spindle-, or mitosis-related GO terms (Fig. S3D and Table S2).

Because the most distinct cellular feature of LP-hESCs other than the survival advantage (12) was mitotic aberration (Fig. 1), the enrichment of the gene signatures of LP-hESCs in the “microtubule organization,” “mitotic spindle organization,” or “cell cycle G2M transition” categories was intriguing (Fig. 3F). Accordingly, the microtubule organization would be different between LP-hESCs and P1 hESCs. To examine this idea, the microtubule stability of P3 hESCs relative to P1 hESCs was determined using the cold treatment protocol (30). As predicted, the mitotic spindle in P3 hESCs remained stabilized compared to P1 hESCs’ mitotic spindles (Fig. 3G). Importantly, microtubule dynamics (e.g., spatiotemporal polymerization and depolymerization) are critical for proper chromosome segregation. Thus, gain of 20q11.21 and following alteration of gene expressions, would be expected to significantly influence microtubule dynamics, thus providing an explanation for the high incidence of CIN in LP-hESCs.

### Prediction of TPX2 for aberrant mitosis in long-term cultured hESCs

Among the genes within 20q11.21 (Fig. 3D), we attempted to identify a gene responsible for aberrant mitosis and microtubule organization or stability, which were both observed in LP-hESCs (Figs. 1 and 3G). Chromosome mis-segregation (Fig. 1B) may result from altered microtubule dynamics, whereas multipolar spindles (Fig. 1E) are possibly due to robust nucleation and the formation of ectopic spindle poles (35). Interestingly, among the genes at the 20q11.21 locus, *TPX2* and *KIF3B* were consistently associated with GO terms such as “microtubule cytoskeleton organization”, “spindle organization”, and “mitotic nuclear division” (Fig. S3D). More importantly, *TPX2,* which encodes a multifunctional protein that has diverse roles in microtubule nucleation and spindle assembly(25) is closely associated with the highest CIN score among multiple genes associated with cancer prognosis (29). Notably, *TPX2* was the gene that contributed most to the enrichment of GO terms of the “spindle pole” and “cell cycle G2M phase transition” categories in GSEA results (Fig. S3D). Therefore, we hypothesized that the upregulation of *TPX2* in LP-hESCs would be accountable for chromosomal abnormalities resulting from perturbed microtubule dynamics.

### Upregulation of TPX2 stabilizes mitotic spindle

To determine the genomic levels of *TPX2* in LP-hESCs, we analyzed genome expression patterns using a whole-genome SNP array. As expected, the fluorescence intensity of genetic probes bound to *TPX2* gradually increased in P3 and P4 (Fig. 4A), and the *TPX2* mRNA levels were also clearly upregulated in P3 and P4 hESCs (Fig. 4B). Consistent with the increased *TPX2* mRNA levels, protein levels were also markedly upregulated in P3 and P4 hESCs (Fig. 4C). It is noteworthy that TPX2 serves as a positive regulator of Aurora A kinase (encoded by *AURKA*) (26), activity of which is also critical for not only microtubule dynamics (36) but also the maintenance of the pluripotency of ESCs through its suppression of the p53 response (37). Accordingly, active phosphorylation of Aurora A was evident in P3 and P4 hESCs along with high TPX2 expression (Fig. 4C). Because there were no noticeable changes to the genomic and mRNA levels of *AURKA* (Fig. S4A) as well as a comparable cell cycle profile (Fig. S4B), the high activity of Aurora A in P3 and P4 hESCs could primarily be attributed to the high expression of TPX2 in P3 and P4 hESCs (Fig. 4C).

**Figure 4.**
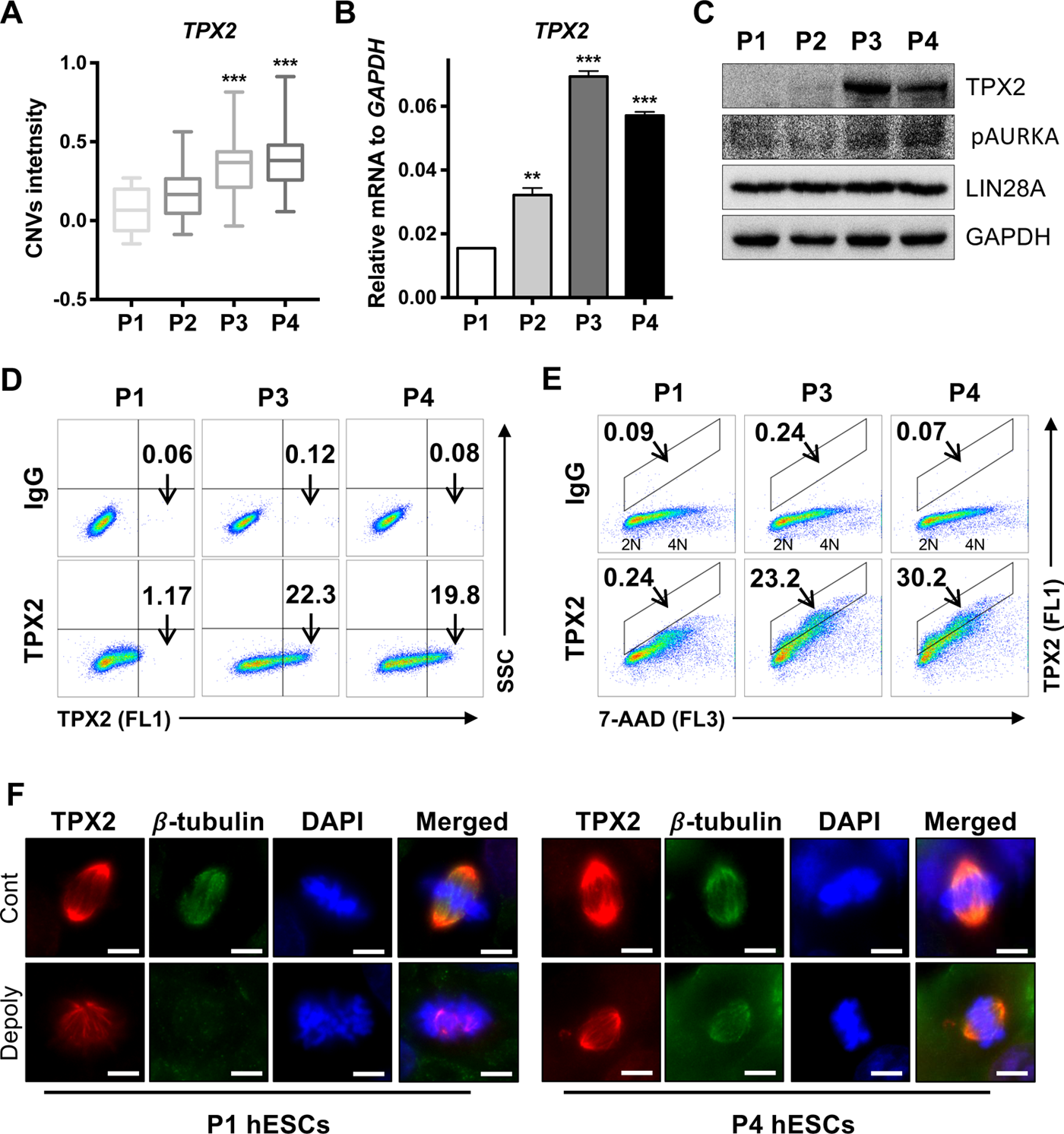
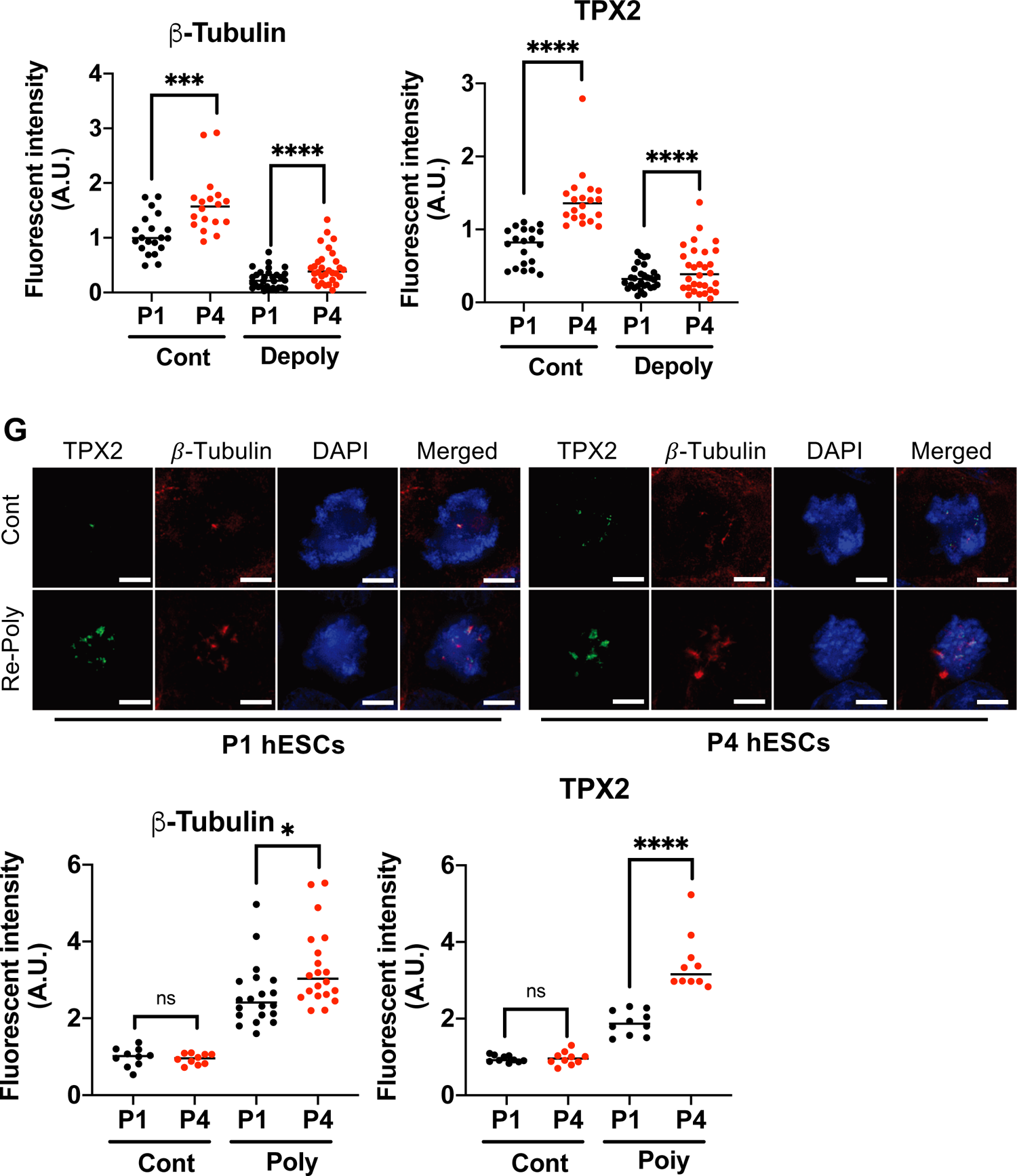
Upregulation of TPX2 stabilizes mitotic spindle. (A-C) Intensity of genetic probes (A), mRNA expression (B), or protein levels (C) for TPX2 were assessed by SNP array (mean ± SD; n = 16, \*\*\**p* < 0.001), real-time PCR (mean ± SD; n = 3, \*\**p* < 0.01 and \*\*\**p* < 0.001), or immunoblotting, respectively. LIN28A or GAPDH was used as a pluripotency marker or a loading control, respectively. (D-E) Flow cytometry analysis for TPX2 (D) or TPX2 co-stained with 7-AAD for DNA contents (E), Arrow indicates the percentages of TPX2^High^ population. (F) Fluorescent microscopic images of mitotic spindle stained with β-tubulin antibody (β-tubulin) and TPX2 at 3 minutes after 1μg/ml of Nocodazole (Noc) treatment (Depoly) in P1 and P4 hESCs (top), Graphical presentation of fluorescent intensity of β-tubulin (left) and TPX2 (right) (scale bar = 10µm) (mean ± SD; n > 16, *** *p*<0.001 and **** *p*<0.0001) (G) Fluorescent microscopic images of mitotic spindle stained with β-tubulin antibody (β-tubulin) and TPX2 at 3 minutes after wash-off of 1μg/ml of Nocodazole (Noc) treatment (Poly) in P1 and P4 hESCs (top), Graphical presentation of fluorescent intensity of β-tubulin (left) and TPX2 (right) (scale bar = 10µm) (mean ± SD; n > 10, ns: not significant, \**p* < 0.05, \*\**p* < 0.01, *** *p*<0.001 and **** *p*<0.0001)

Of special interest, roughly 20% of P3 and P4 hESC populations highly expressed TPX2. (Fig. 4D). Although the TPX2 protein, which is regulated in a cell cycle-dependent manner, is highly upregulated in the G2/M phase(25), the high TPX2-positive populations in P3 and P4 cells would not likely have resulted from a larger proportion of G2/M cells among P3 and P4 hESCs, as these cells exhibited similar cell cycle profiles compared to P1 hESCs (Figs. S1E and S4B). To compare expression levels among these hESC populations, TPX2 expression was determined by dual staining of TPX2 and DNA content with 7-AAD (Fig. 4E). The similar cell cycle profiles (Figs. S1E and S4B) and TPX2-positive signals, along with DNA content, (Fig. 4E) of P3 and P4 hESCs compared to P1 hESCs clearly reveal that TPX2 expression is generally upregulated regardless of cell cycle (Fig. 4E). Furthermore, mitotic spindle integrity (determined by β-tubulin fluorescence intensity), which is strongly associated with TPX2 expression, was clearly stabilized in P4 hESCs compared to P1 hESCs after depolymerization was induced with a nocodazole treatment (Fig. 4F). Repolymerization of the mitotic spindle after release from the nocodazole treatment was also facilitated in P4 hESCs along with spindle arrangement by TPX2 (Fig. 4G).

### Establishment of inducible TPX2 in early passage hESCs

To examine the effects of high TPX2 expression on microtubule dynamics and consequent mitotic event (Fig. 1), we attempted to stably express TPX2 in P1 hESCs. However, for unknown reasons, we failed to stably express TPX2 in hESCs even after multiple trials (data not shown). Instead, TPX2 expression was conditionally expressed through a doxycycline (Dox) inducible system in P1 hESCs (iTPX2 hESCs) (Fig. 5A). After Dox treatment, the clear induction of mRNA and the production of TPX2 protein was observed (Fig. 5B), along with fluorescence of an enhanced green fluorescent protein (eGFP) tagged to TPX2 (Fig. 5C). The eGFP signal from the mitotic cells after Dox treatment was clearly associated with the mitotic spindle where the mitotic TPX2 is localized (Fig. 5D). We noticed that the prolonged expression of TPX2 by Dox treatment suppressed cell proliferation (Fig. 5E), which would account for the repeated failure to establish TPX2-stable hESCs. Based on the observed increase in the 4N peak in the eGFP-positive population after Dox treatment (Fig. 5F), the growth suppression by stable expression of TPX2 resulted from the induction of G2/M arrest. TPX2 induction (10-15-fold compared to P1 hESCs) by Dox (1 μg/mL) resulted in G2/M arrest and growth suppression in iTPX2 hESCs (Fig. 5G). Of note, the basal level of TPX2 in iTPX2 hESCs was comparable to that of P4 hESCs; this may have resulted from the leaky expression of the Tet-On system (38) (Fig. 5G). We then aimed to determine the optimal dose of Dox to induce a moderate level of TPX2 that may not interfere with growth. TPX2 induction in iTPX2 hESCs occurred at Dox concentrations as low as 0.05 μg/mL and dramatically increased in a dose-dependent manner (Fig. 5H). Colony formation at various doses of Dox revealed that hESC growth was minimally affected at a dosage of 0.1 μg/mL (Fig. 5I), where moderate levels of TPX2 were achieved (Fig. 5H).

**Figure 5.**
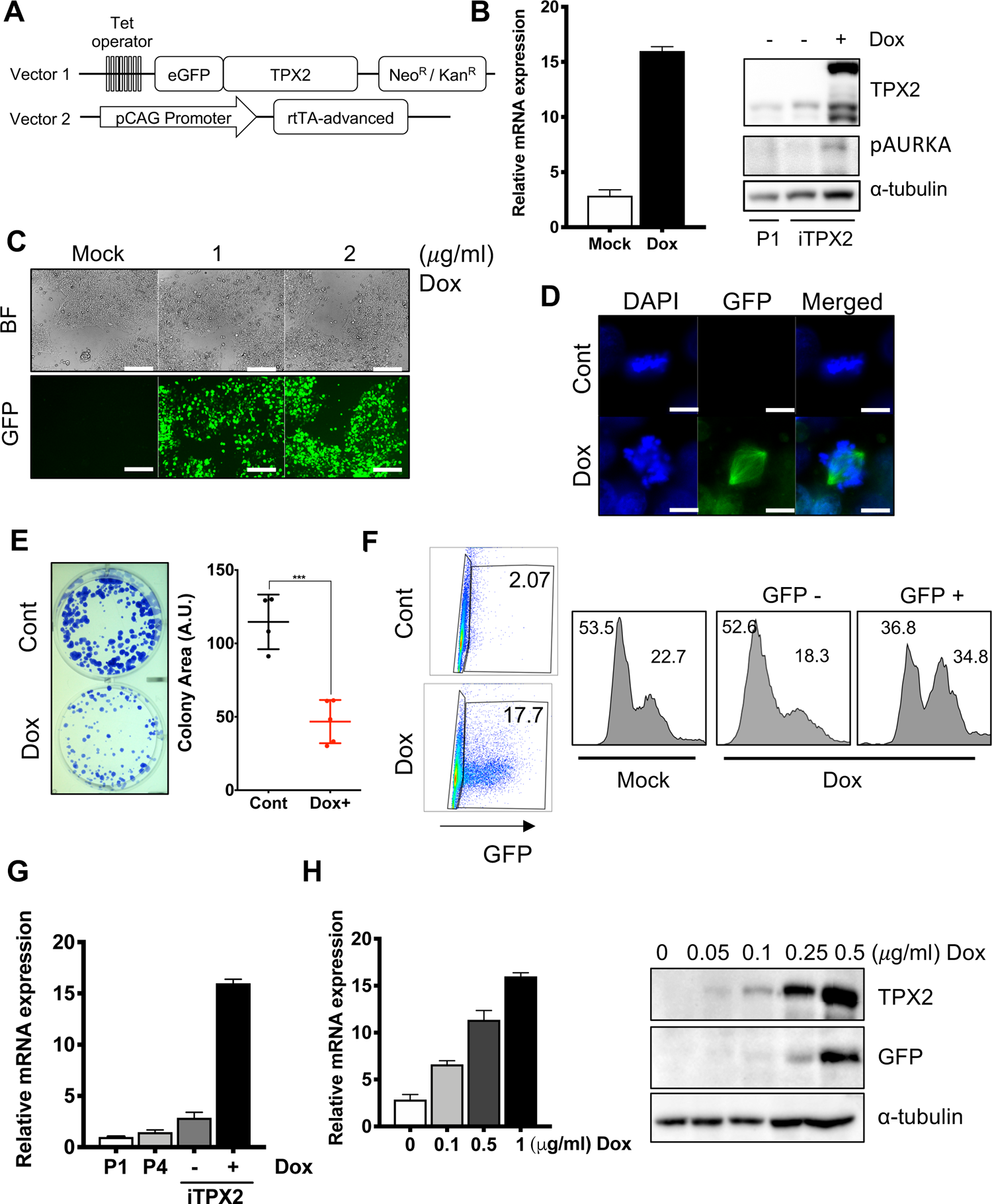
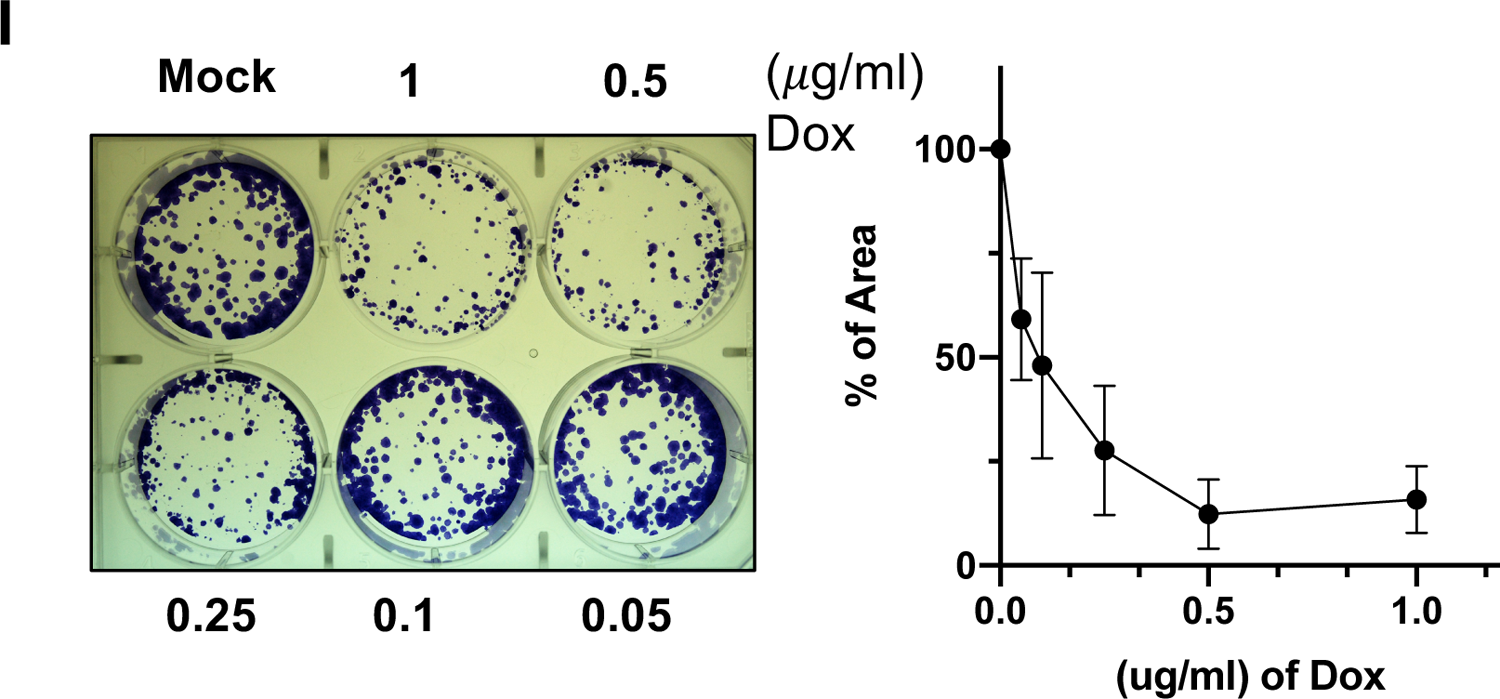
Establishment of inducible TPX2 in early passage hESCs. (A) Schematic presentation of the vector used for establishment of EGFP-tagged doxycycline inducible TPX2 (iTPX2) hESCs (B) Relative mRNA expression (left) and protein (right) level of TPX2 in iTPX2 hESCs in 24 hours after 1μg/ml of Dox treatment compared to P1 hESCs (P1) (C) Light (top) and fluorescent (bottom) microscopic images of iTPX2 hESCs in 24 hours after indicated concentration of Dox treatment (scale bar = 50µm) (D) Fluorescent microscopic images of mitotic cells of iTPX2 hESCs treated with or without 1μg/ml of Dox (scale bar = 10µm) (E) Clonogenic assay of iTPX2 hESCs with or without 1μg/ml of Dox (left) and graphical presentation of number of colony (right) (mean ± SD; n = 5 *** *p*<0.001) (F) Flow cytometry for DNA content of GFP negative (GFP-) and positive (GFP+) of iTPX2 hESCs after 1μg/ml of Dox treatment (G) Relative expression level of TPX2 of P1, P4 hESCs and iTPX2 hESCs with or without 1μg/ml of Dox treatment (n = 2) (H) Expression level of mRNA (left) and protein (right) of TPX2 in iTPX2 hESCs at indicated dose of Dox (n = 2) (I) Clonogenic assay of iTPX2 hESCs after indicated dose of Dox treatment (left) and graphical presentation of number of colony (right) (n = 2)

### Spindle stabilization and abnormal chromosome condensation by TPX2 induction

Next, we examined whether TPX2 expression affects mitosis in hESCs. Using a Dox concentration that exerts only marginal effects on cell growth (Fig. 5I), the duration of mitosis with or without TPX2 expression was assessed. Similar to P4 hESCs compared to P1 hESCs, the mitotic duration of iTPX2 cells was significantly delayed by TPX2 induction (Fig. 6A). It is generally accepted that the disruption of spindle dynamics affects mitotic duration. In line with this, the extended mitotic duration observed as a result of TPX2 expression may occur because of disarrangement of spindle dynamics. To this end, the assembly or disassembly of mitotic spindles was monitored with or without the moderate induction of TPX2 in iTPX2 hESCs. Similar to the high stability of the mitotic spindles of P4 hESCs, where TPX2 expression was higher than P1 hESCs (Figs. 4F and G), Dox-induced TPX2 expression in iTPX2 hESCs stabilized the mitotic spindle upon depolymerization (Fig. 6B). While the mitotic spindle was completely depolymerized at 9 minutes after nocodazole treatment, in the iTPX2 where TPX2 expression was induced by Dox, the spindle remained partly polymerized (Fig. 6C). Consistent repolymerization of the mitotic spindle after the removal of nocodazole was significantly enhanced by Dox treatment (Fig. 6D). This data clearly demonstrates that high TPX2 expression stabilizes the mitotic spindle or promotes spindle assembly, leading to prolonged mitotic duration. Next, to closely monitor the mitotic cells expressing high levels of TPX2, individual mitotic cells were examined by immunofluorescence. Surprisingly, we observed widespread mitotic chromosome condensation abnormalities in iTPX2 hESCs that had undergone Dox treatment for 48 hours. Unlike P1 hESCs or iTPX2 hESCs which did not undergo Dox treatment, chromosome alignment failed even in metaphase (Fig. 6E). In particular, typical spindle fibers depolymerization observed at telophase of P1 was largely disorganized at iTPX2 hESCs by TPX2 induction (Fig 6F). These evident disorders occurring in mitotic spindles would be strongly associated to drastic event of polyploid in iTPX2 hESCs after TPX2 induction (Fig. 6G). Overall, our findings indicate that the high expression of TPX2 observed in LP-hESCs would delay mitotic progression and induce polyploidy by stabilizing the mitotic spindle.

**Figure 6.**
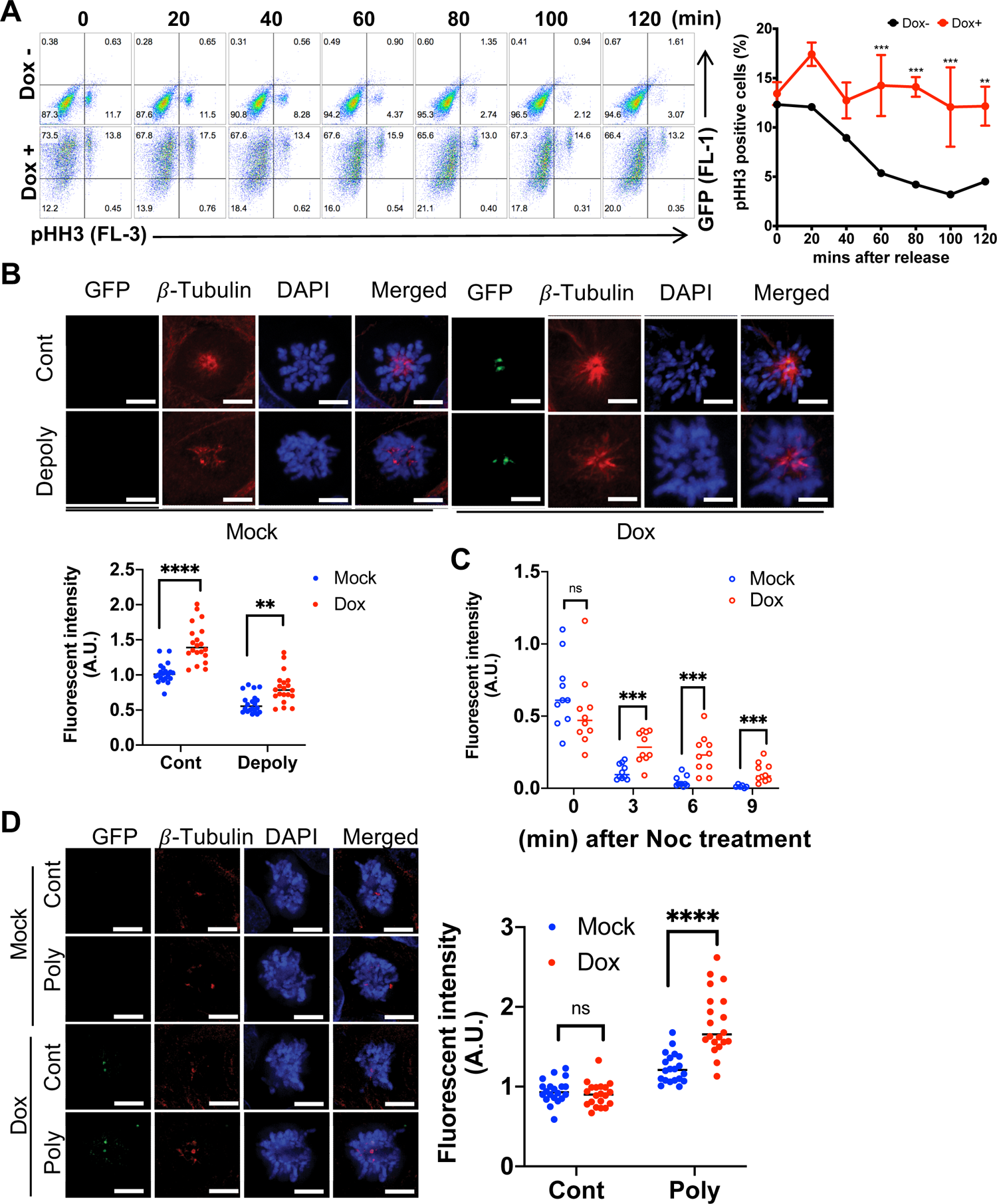
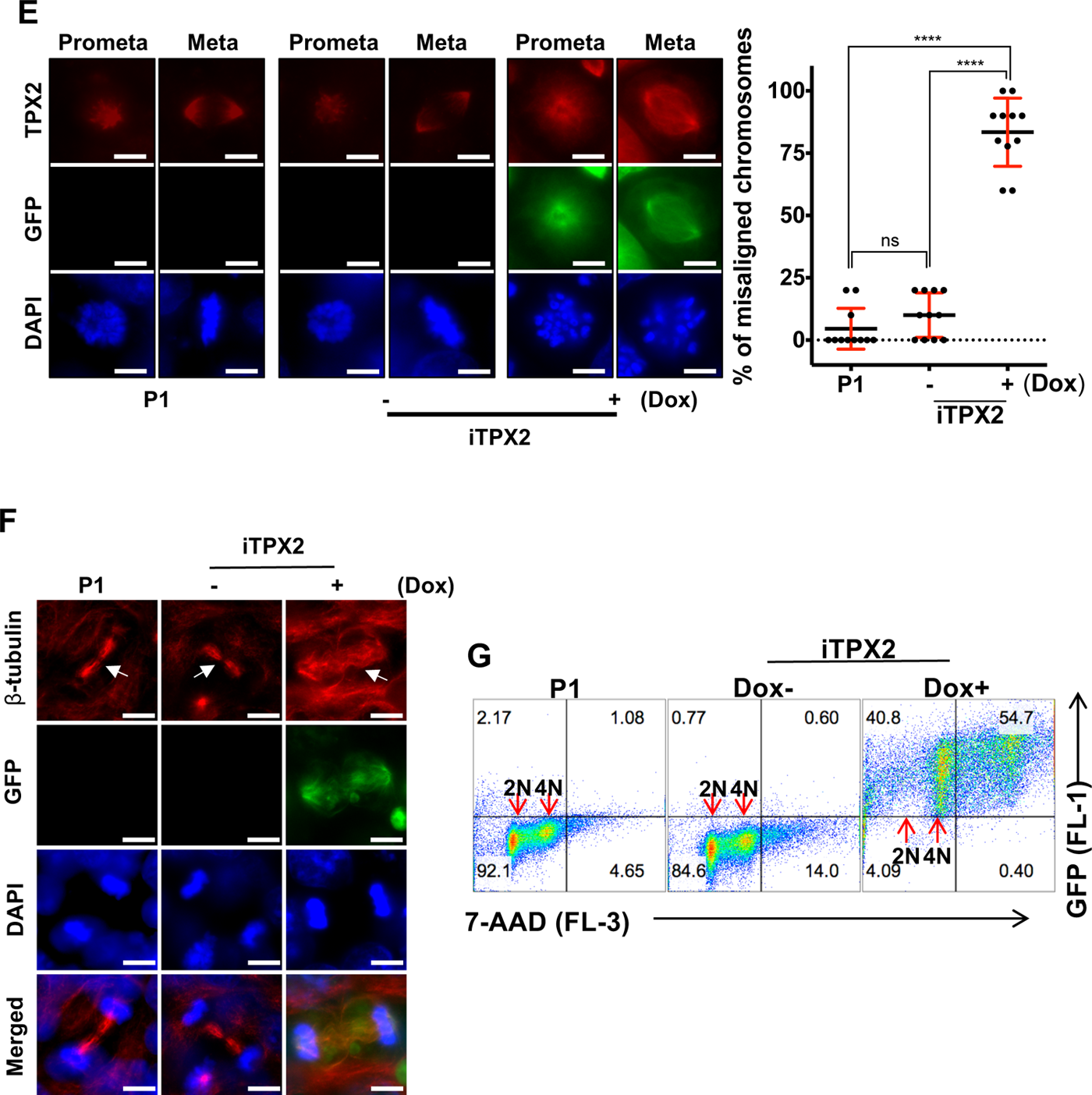
Spindle stabilization and abnormal chromosome condensate by TPX2 induction. (A) Flow-cytometry for mitotic population of iTPX2 hESCs with or without Dox upon release from mitotic arrest with nocodazole (25ng/mL) (left), Graphical presentation of Phospho-histone H3 (pHH3) positive cells with or without Dox treatment, at indicated time after release (right) (mean ± SD; n = 2, \*\**p* < 0.01, *** *p*<0.001 and **** *p*<0.0001) (B) Florescent images of mitotic spindle stained with β tubulin antibody and GFP of pro-metaphase iTPX2 hESCs with (Dox) or without (Mock) Dox treatment before (Cont) and 3 minutes after 1μg/ml of nocodazole (Depoly) (top), Graphical presentation of fluorescent intensity of β-tubulin (bottom) (scale bar = 10µm) (mean ± SD; n > 15, \*\**p* < 0.01, *** *p*<0.001 and **** *p*<0.0001) (C) Graphical presentation of fluorescent intensity of β-tubulin at indicated time after 1μg/ml of nocodazole (mean ± SD; n > 9, ns: not significant, *** *p*<0.001) (D) Florescent images of mitotic spindle stained with β–tubulin antibody and GFP of pro-metaphase iTPX2 hESCs with (Dox) or without (Mock) Dox treatment before (Cont) and 5 minutes after wash-off of 1μg/ml of nocodazole (Poly) (left), Graphical presentation of fluorescent intensity of β-tubulin (right) (scale bar = 10µm) (mean ± SD; n > 15, ns: not significant, **** *p*<0.0001) (E) Florescent images of prometaphase (Prometa) and metaphase (Meta) cells of P1 and iTPX2 hESCs at 48 hours after Dox (left), Graphical presentation of percentage of misaligned mitotic chromosome (right) (scale bar = 10µm) (mean ± SD; n = 11, ns: not significant, **** *p*<0.0001) (F) Florescent images of mitotic spindle stained with β antibody (indicated by white arrow) and GFP in telophase of P1 and iTPX2 hESCs at 48hrs after Dox treatment (scale bar = 10µm) (G) Flow-cytometry of DNA content of P1 and iTPX2 hESCs at 48hrs after Dox treatment (top), Graphical presentation of DNA content of indicated condition (bottom) (H) CGH array result (ns: not significant, \**p* < 0.05, \*\**p* < 0.01, *** *p*<0.001 and **** *p*<0.0001)

## Discussion

In this study, we found that culture-adapted hESCs acquire a functional signature associated with a gain of 20q11.21 function. GSEA revealed these cells to be highly enriched in the biological process for “microtubule dynamics at G2/M.” By multiple approaches, we identified *TPX2*, located in 20q11.21, as a putative driver for chromosome misalignment and consequent mitotic abnormality, as inducible expression of TPX2 in EP-hESCs affected mitotic progression and spindle dynamics, which were similarly observed in LP-hESCs.

The maintenance of genome integrity in human pluripotent stem cells (hPSCs) is exhaustively challenged by simultaneous exposure to multiple artificial selective pressures during *in vitro* propagation. Given that hPSCs are vulnerable to genomic insults (1), they progressively acquire genetic alterations or mutations upon prolonged culturing, as shown in Figure 2. Thereafter, those clones that survive, are able to do so because they have acquired either resistance to apoptosis or rapid proliferation rates (20), thereby becoming predominant in the cultures (Figs. 1F and S1B). These aberrant hPSCs exhibit recurrent CNVs at chromosomes 12, 17, 20, and X (3, 21). The amplification of sub-chromosomal 20q11.21 has emerged as a frequent variant in culture-adapted hPSCs (9), a discovery propelled by advances in single-nucleotide resolution of genome-wide approaches. Considering that 25% of normal karyotype hESC lines, including hPSCs with early passage (14), and our cell models (Figs. 2 and 3) revealed a gain of 20q11.21 (9), it is likely hard to detect this minimal amplicon using conventional cytogenetic testing methods (i.e., spectral karyotyping and G-band karyotyping). This suggests that genetically altered hPSCs possessing malignancy or chemoresistance acquired by gaining this region cannot be readily perceived (11). Notably, the gain of 20q11.21 has only been detected in cultured hPSCs and not during embryo derivation (10).

Indeed, our results demonstrate that culture-adapted hESCs are resistant to apoptosis (Fig. S1C) and consequently possess survival advantages in cultures (Figs. 1F and S1B). More interestingly, chromosome misalignment, aberrant mitosis (Figs. 1A-E), and consequent polyploidy (Fig. 1G) were frequently observed in LP-hESCs in our study, which is consistent with previous reports (39, 40). Although the amplification of 20q11.21 has been determined to provide growth advantages mediated by increased levels of anti-apoptotic factor *BCL2L1* (11), there is no clear evidence that *BCL2L1* is involved in the molecular mechanisms underlying the incidence of CIN. Therefore, to uncover the biological cause of CIN in LP-hESCs, we performed GO functional analysis for genes commonly amplified in both CNVs and the gene expression profiles of LP-hESCs (Fig. 3). Surprisingly, the genes were almost exclusively included in the 20q11.21 region and were highly enriched in spindle/microtubule or G2M transition annotation (Fig. 3F), a process responsible for maintenance of chromosome integrity (41). Thus, these results indicate that gaining 20q11.21 can confer survival advantages and chromosomal abnormalities in culture-adapted hPSCs.

Given that *TPX2* is commonly associated with the “microtubule regulation” GO term and the 20q11.21 region (Fig. S3D), the gain of 20q11.21-driven *TPX2* amplification would likely be responsible for the deregulation of microtubule organization, which can further lead to chromosome misalignment and abnormal mitosis (29). Indeed, as shown in Figure 4, both the mRNA and protein levels of TPX2 were significantly upregulated in LP-hESCs, which corresponded with high microtubule stability (Fig. 4). TPX2 has frequently been reported to be overexpressed in various tumor types (25), acting as an oncogenic holoenzyme with Aurora kinase A (AURKA) by engaging its enzymatic activity (26) to be a potential target for genomically unstable cancer cells (42). As expected, TPX2 induction in P1-hESCs activated Aurora A kinase (Fig. 6B) and markedly affected mitotic spindle dynamics (Fig. 7), which was similar our observations in P4-hESCs (Figs. 1A and 4C). Thus, our findings indicate that *TPX2* amplification triggers chromosome misalignment and aberrant mitosis by controlling microtubule kinetics. It is noteworthy that *TPX2* overexpression alone failed to result in cellular transformation but not abnormal spindle formation (26).

In conclusion, when hPSCs acquire survival advantages by gaining 20q11.21 during prolonged culturing, concurrent *TPX2* amplification leads to sustained microtubule stabilization, which consequently causes chromosomal abnormalities in the culture-selected hPSCs.

## Author contributions

HJ.C conceived the overall study design, led the experiments and wrote the manuscript. HC.J and HY.G conducted the experiments, data analysis and wrote the first draft. JG.S and HD.S performed CGH array and exome sequencing and analyzed the data. YJ.K validated the experiments and generated cell lines. MG.C, D.G, HS.C, JH.L and CY.J performed cell cycle analysis and measured the spindle stability. All authors contributed to manuscript writing and revising and endorsed the final version of this manuscript.

## Acknowledgments

We thank Dr. Minh Dang Nguyen in University of Calgary for kindly providing TPX2 constructs. This work was supported by a grant from the National Research Foundation of Korea (NRF) (grant number: 2020M3A9E4037905 and 2020R1A2C2005914).

