## Supplemental Materials for "TPX2 Amplification-Driven Aberrant Mitosis in Long-Term Cultured Human Embryonic Stem Cells"

This PDF file includes:

Supplemental Material and Methods

Supplemental Figure Legends

**Supplemental Material and Methods**

**Live cell imaging**

Cells were placed on 4-well Chambered Coverglass (Thermo Scientific) coated with matrigel and then incubated with 1 μg/ml of Hoechst 33342 (Thermo, H1399) to visualize chromosomes. Fluorescent images were acquired every 5 min for 24 hrs using a Nikon eclipse Ti with a 40x dry Plan-Apochromat objective, captured with an iXonEM +897 Electron Multiplying charge-coupled device camera, and analyzed using a NIS elements Ar microscope imaging software.

**Immunoblotting and Immunofluorescence**

Cells were lysed with tissue lysis buffer (TLB, 20mM Tric-HCl, pH 7.4, 137mM NaCl, 2mM EDTA, 1% Triton X-100, and 10% glycerol) supplemented with 0.2mM sodium vanadate and 1mM protease inhibitor cocktail (Roche), and Immunoblotting assay was performed as described previously [65]. Primary antibodies used in this study were as followed: TPX2 (Cell signaling, #12245), pAurora A (Cell signaling, #3079), Aurora A (Cell signaling, #4718), LIN28A (Cell signaling, #3979), GAPDH (AbFrontier), Cyclin B1 (Santa cruz, sc-245), and α-tubulin (Santa cruz, sc-8035). For Immunofluorescence, cells were fixed with 4% paraformaldehyde, permeabilized with 0.25% Triton X-100, and then blocked with TBS-T containing 3% BSA. The coverslips were incubated with primary antibodies: α-tubulin (1:400), β-tubulin (1:400), and Aurora A (1:200) at 4°C overnight and stained with either Cy2- (Jackson ImmunoResearch, 1:200) or Alexa 594- (Life Technologies, 1:200) conjugated secondary antibody with DAPI (1:200) at room temperature for 1 hr. Cell images were captured and analyzed by a BX53 research microscope.

**Flow cytometry analysis**

1 x $10^6$ cells were fixed/permeabilized with BD Cytofix/Cytoperm (BD Biosciences, 554722) and stained with following primary antibodies: Oct4 (Abcam, ab19857, 1:100), Nanog (Cell signaling, #4903, 1:100), and TPX2 (Cell signaling, 1:200), followed by incubation with Alexa 488- (Life Technologies, 1:200) or PerCP- (R&D, F0114, 10µl per each sample) conjugated secondary antibodies. The cells were washed, suspended in BD Perm/Wash (BD Biosciences, 554723), and measured by flow cytometry on a FACSCalibur (BD Biosciences). For SSEA4 staining, fluorescence-labeled SSEA4-FITC antibodies (BD Pharmingen, 560126) were used. To detect DNA contents, cells were fixed with chilled 70% ethanol at 4°C overnight, incubated with RNase A (Sigma, 100µg/ml) and propidium iodide (Sigma, 50µg/ml) in dark room, and analyzed by the flow cytometer. The acquired data were analyzed by FlowJo software.

**Fluorescence-based competitive proliferation assay**

GFP-expressing hESC (EGFP-P1) and P3 hESC are cultured together. Cells are detached with Accutase (BD Bioscience) and rinsed with DPBS three times before flow cytometry. GFP+ cells in the total population are measured using flow cytometry.

#### Primer information

**Table 1**. Primer Sequences for quantitative real-time PCR analysis

| Gene | Forward Sequence (5' to 3') | Reverse Sequence (5' to 3') |
|---|---|---|
| ***TPX2*** | GCTCAACCTGTGCCACATTA | CGAGAAAGGGCATATTTCCA |
| ***AURKA*** | TCCTGAGGAGGAACTGGCATCAAA | TACCCAGAGGGCGACCAATTTCAA |

#### Supplemental Figure Legends

**Figure S1.** (A) Short tandem repeat profile of P1, P3, or P4 hESCs compared to that of WiCell H9 hESC line (B) Competitive cell growth assay for P3 hESCs mixed with P1 hESCs expressing EGFP (C) Flow cytometry analysis for Annexin V and 7-AAD, Cells were treated with etoposide for 24 hrs and the percentages of live cells (dual negative for Annexin V and 7-AAD) were presented. (D) Flow cytometry analysis for pluripotency markers in P1, P3, or P4 hESCs (E) BrdU incorporation assay was performed in P1, P3, or P4 hESCs. The percentages of active proliferating cells (BrdU+) were graphically represented in lower right panel.

**Figure S2.** (A) Log R ratio (LRR) plots for chromosome 17 region assessed by SNP array, Individual intensity of genetic probes is represented as a blue-filled circle. (B) The LRR plots for 17q24 (about 3 Mb sub-chromosomal region) were magnified and the y-axis ranges from -2 to 2.

**Figure S3.** (A) Overall gene expression patterns of the four hESC lines (P1-P4) in duplicate. TPM was taken as gene expression level for each sample. A total of 11,203 genes with standard deviation > 0.1 were used. (B) Volcano plots showing DEGs in P2, P3, and P4 compared to P1 hESCs, and DEGs in LP-hESCs compared to EP-hESCs. DEGs were selected using DESeq2 with $|\log_2\text{fold-change}|>1$ and a false discovery rate (FDR) < 0.01. (C) Heatmap showing gene set similarity of significantly enriched GO Biological Process (BP) terms in LP-hESCs compared to EP-hESCs. Similarity score was driven by the hypergeometric *P*-value. Representative GO terms of each cluster were marked as underline. (D) GSEA results for microtubule-related GO terms. The top 5 leading genes were selected from the leading-edge subsets, defined as the genes that contributed the most to the enrichment signal of a given set.

**Figure S4.** (A) CNV intensity for *AURKA* was assessed by SNP array. (B) Cell cycle profiles in P1, P3, or P4 hESCs, DNA contents of cells were analyzed by PI staining.

**Movie S1 Time lapse images of P1 (A), P2 (B), P3 (C) and P4 (D) hESCs at mitosis**

**Table S1 List of differentiated expressed genes between EP and LP-hESCs**

**Table S2 List of upregulated functions in LP-hESCs**

### Figure. S1

## A

| Samples | D16S539 | D7S820 | D13S317 | D5S818 | CSF1PO | TPOX | Amelogenin | TH01 | vWA |
|---|---|---|---|---|---|---|---|---|---|
| H9 (WiCell) | **12, 13** | **9, 11** | **9, 9** | **11, 12** | **11, 11** | **10, 11** | **X, X** | **9.3, 9.3** | **17, 17** |
| P1 (40s) | **12, 13** | **9, 11** | **9, 9** | **11, 12** | **11, 11** | **10, 11** | **X, X** | **9.3, 9.3** | **17, 17** |
| P3 (200s) | **12, 13** | **9, 11** | **9, 9** | **11, 12** | **11, 11** | **10, 11** | **X, X** | **9.3, 9.3** | **17, 17** |
| P4 (300s) | **12, 13** | **9, 11** | **9, 9** | **11, 12** | **11, 11** | **10, 11** | **X, X** | **9.3, 9.3** | **17, 17** |

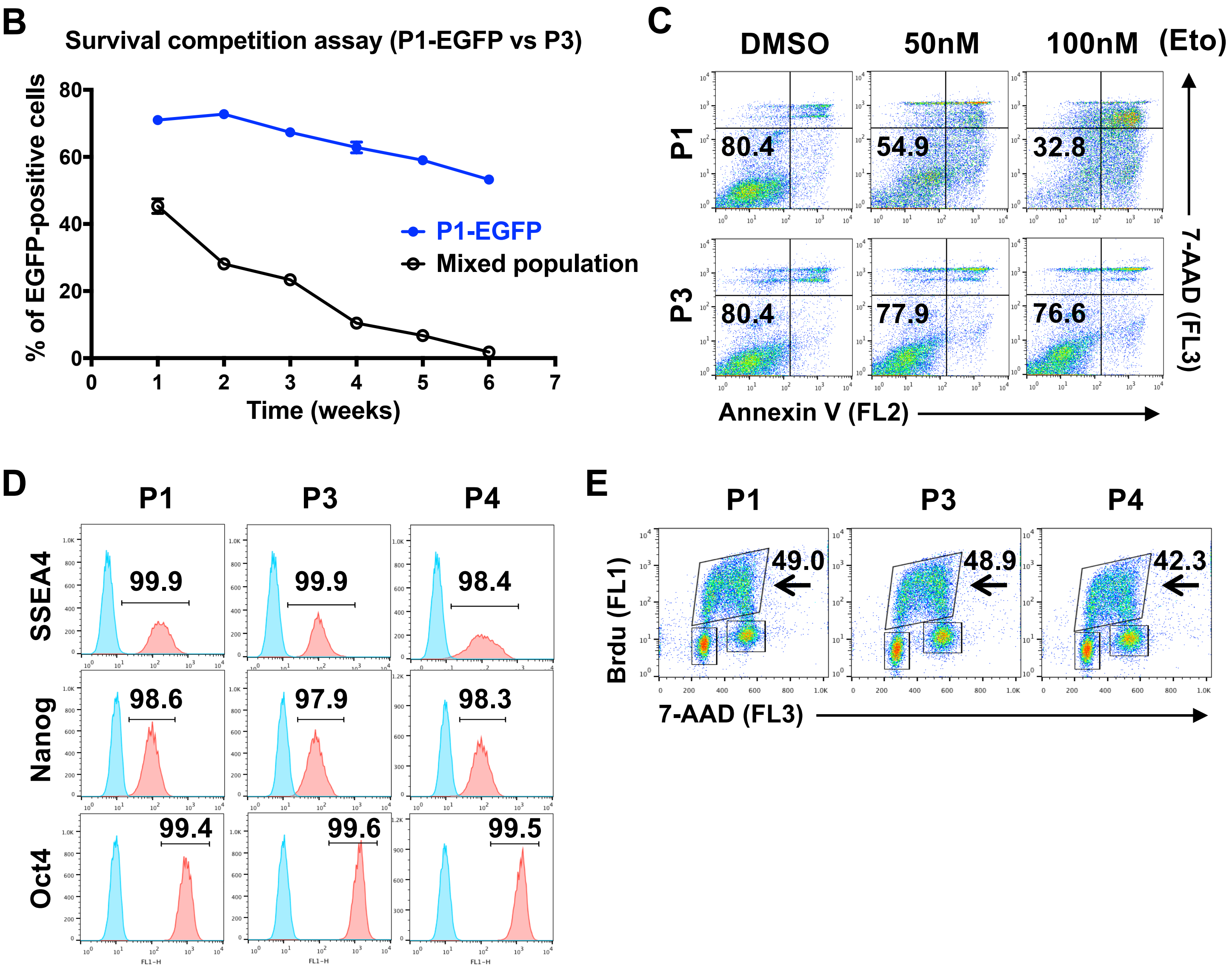

### Figure. S2

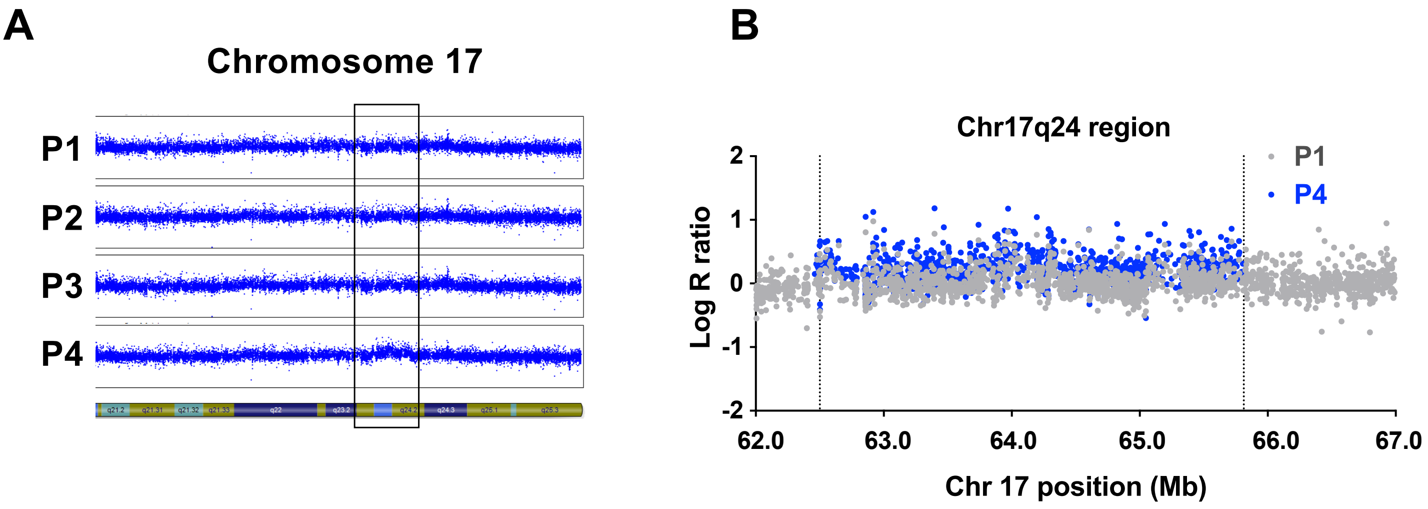
A
Chromosome 17
P1
P2
P3
P4
B
Chr17q24 region
P1
P4
Log R ratio
2
1
0
-1
-2
62.0
63.0
64.0
65.0
66.0
67.0
Chr 17 position (Mb)

### Figure. S3

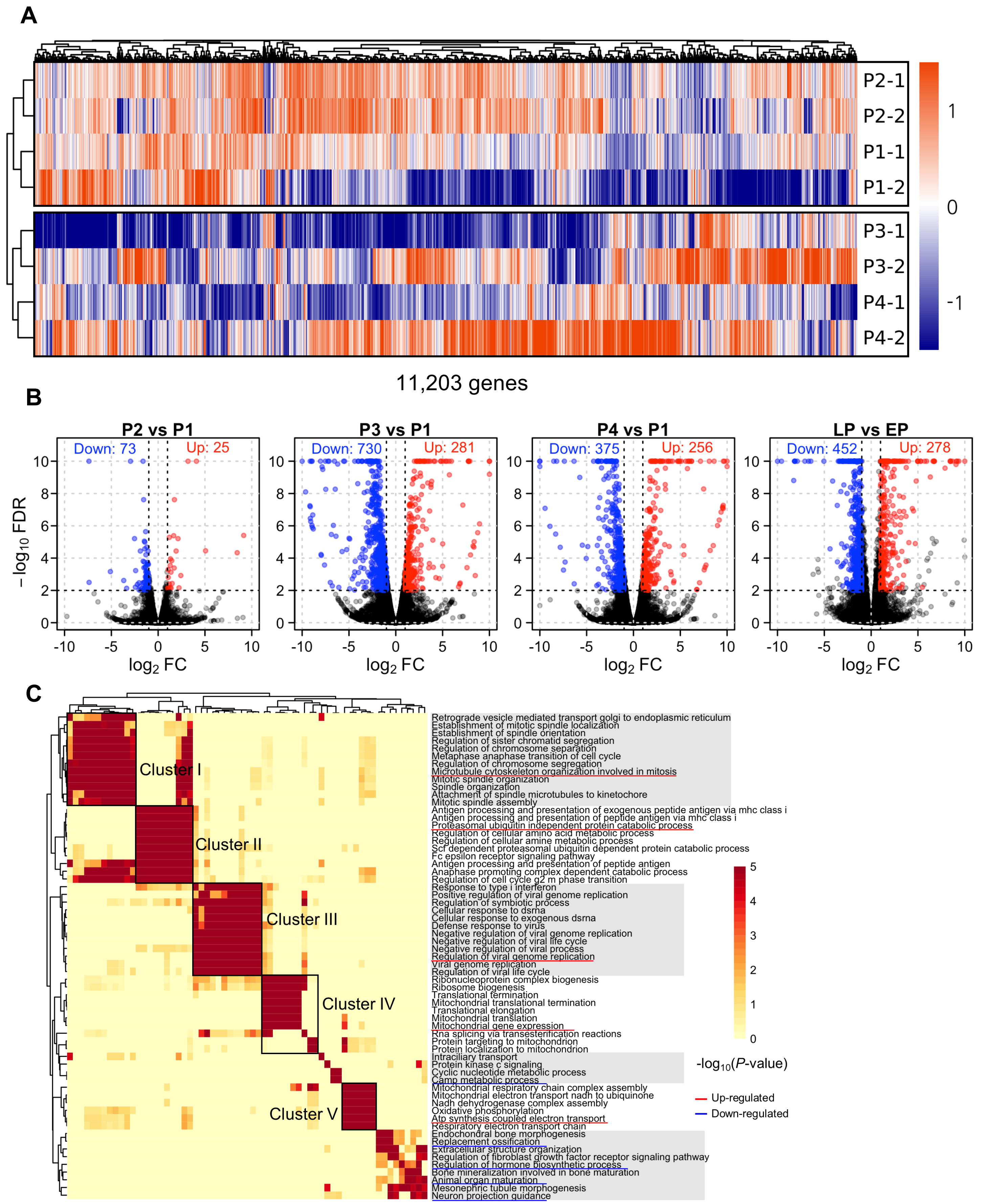

A
P2-1
P2-2
P1-1
P1-2
P3-1
P3-2
P4-1
P4-2
1
0
-1
11,203 genes
B
P2 vs P1
Down: 73
Up: 25
P3 vs P1
Down: 730
Up: 281
P4 vs P1
Down: 375
Up: 256
LP vs EP
Down: 452
Up: 278
$-\log_{10}$ FDR
$\log_2$ FC
C
Cluster I
Cluster II
Cluster III
Cluster IV
Cluster V
Retrograde vesicle mediated transport golgi to endoplasmic reticulum
Establishment of mitotic spindle localization
Establishment of spindle orientation
Regulation of sister chromatid segregation
Regulation of chromosome separation
Metaphase anaphase transition of cell cycle
Regulation of chromosome segregation
Microtubule cytoskeleton organization involved in mitosis
Mitotic spindle organization
Spindle organization
Attachment of spindle microtubules to kinetochore
Mitotic spindle assembly
Antigen processing and presentation of exogenous peptide antigen via mhc class i
Antigen processing and presentation of peptide antigen via mhc class i
Proteasomal ubiquitin independent protein catabolic process
Regulation of cellular amino acid metabolic process
Regulation of cellular amine metabolic process
Scf dependent proteasomal ubiquitin dependent protein catabolic process
Fc epsilon receptor signaling pathway
Antigen processing and presentation of peptide antigen
Anaphase promoting complex dependent catabolic process
Regulation of cell cycle g2 m phase transition
Response to type i interferon
Positive regulation of viral genome replication
Regulation of symbiotic process
Cellular response to dsrna
Cellular response to exogenous dsrna
Defense response to virus
Negative regulation of viral genome replication
Negative regulation of viral life cycle
Negative regulation of viral process
Regulation of viral genome replication
Viral genome replication
Regulation of viral life cycle
Ribonucleoprotein complex biogenesis
Ribosome biogenesis
Translational termination
Mitochondrial translational termination
Translational elongation
Mitochondrial translation
Mitochondrial gene expression
Rna splicing via transesterification reactions
Protein targeting to mitochondrion
Protein localization to mitochondrion
Intraciliary transport
Protein kinase c signaling
Cyclic nucleotide metabolic process
Camp metabolic process
Mitochondrial respiratory chain complex assembly
Mitochondrial electron transport nadh to ubiquinone
Nadh dehydrogenase complex assembly
Oxidative phosphorylation
Atp synthesis coupled electron transport
Respiratory electron transport chain
Endochondral bone morphogenesis
Replacement ossification
Extracellular structure organization
Regulation of fibroblast growth factor receptor signaling pathway
Regulation of hormone biosynthetic process
Bone mineralization involved in bone maturation
Animal organ maturation
Mesonephric tubule morphogenesis
Neuron projection guidance
$-\log_{10}(P\text{-value})$
Up-regulated
Down-regulated

D

#### Microtubule, spindle, mitosis related GO terms

| GO term | NES | P-value | Top5 leading genes |
|---|---|---|---|
| Microtubule cytoskeleton organization involved in mitosis | 2.35 | 0.00022 | **KIF3B, TPX2**, AURKC, INSC, NDC80 |
| Mitotic spindle organization | 2.20 | 0.00022 | **KIF3B, TPX2**, AURKC, NDC80, CENPE |
| Establishment of mitotic spindle localization | 2.17 | 0.00021 | INSC, NDC80, ITGB1, MCPH1, ZW10 |
| Attachment of spindle microtubules to kinetochore | 2.08 | 0.00021 | AURKC, NDC80, CENPE, KNSTRN, FAM175B |
| Spindle microtubule | 2.08 | 0.00022 | **KIF3B**, AURKC, CENPE, KIFAP3, KIF2A |
| Establishment of spindle orientation | 2.07 | 0.00021 | INSC, NDC80, ITGB1, MCPH1, ZW10 |
| Mitotic spindle assembly | 2.04 | 0.00021 | **KIF3B, TPX2**, AURKC, FAM175B, KIF2A |
| Spindle organization | 2.02 | 0.00022 | **KIF3B, TPX2**, AURKC, UVRAG, NDC80 |
| Regulation of cell cycle g2 m phase transition | 2.01 | 0.00022 | **TPX2**, OFD1, DYNC1I2, KCNH5, PSMD1 |
| Cell cycle g2 m phase transition | 1.99 | 0.00022 | **TPX2**, OFD1, DYNC1I2, KCNH5, CCNA2 |
| Negative regulation of cell cycle g2 m phase transition | 1.98 | 0.00022 | PSMD1, PSMB1, PSMD12, PSMC6, PSMA1 |
| Spindle assembly | 1.90 | 0.00022 | **KIF3B, TPX2**, AURKC, FAM175B, KIF2A |
| Spindle | 1.85 | 0.00023 | **KIF3B, TPX2**, AURKC, UXT, SNCG |
| Spindle pole | 1.79 | 0.00022 | **TPX2**, AURKC, UXT, BEX4, CKAP2L |
| Microtubule binding | 1.59 | 0.00022 | **KIF3B, TPX2**, JAKMIP2, UXT, MID1 |
| Transport along microtubule | 1.58 | 0.00022 | **KIF3B**, UXT, TRIM58, DYNC1I2, HSPA8 |
| Microtubule | 1.55 | 0.00023 | **KIF3B, TPX2**, AURKC, MID1, BEX4 |
| Spindle localization | 1.85 | 0.00042 | INSC, NDC80, ITGB1, MCPH1, ZW10 |
| Regulation of microtubule cytoskeleton organization | 1.56 | 0.00043 | **TPX2**, MID1, PAK1, FGF13, PHLDB2 |
| Microtubule associated complex | 1.58 | 0.00065 | **KIF3B**, AURKC, MID1, DYNC1I2, PEA15 |
| Microtubule polymerization or depolymerization | 1.68 | 0.00066 | **TPX2**, MID1, PAK1, FGF13, KIF2A |
| Microtubule based transport | 1.52 | 0.00087 | **KIF3B**, UXT, OFD1, TRIM58, DYNC1I2 |
| Microtubule based movement | 1.41 | 0.00092 | **KIF3B**, UXT, OFD1, TRIM58, DYNC1I2 |
| Spindle midzone | 1.88 | 0.00105 | AURKC, CENPE, CDCA8, MAP10, KIF18A |
| Mitotic spindle | 1.59 | 0.00152 | **TPX2**, CENPE, CKAP2L, KNSTRN, MAP10 |
| Spindle midzone assembly | 1.89 | 0.00163 | AURKC, MAP10, RACGAP1, KIF4A, AURKB |
| Regulation of spindle organization | 1.79 | 0.00210 | **TPX2**, DRG1, GNAI1, CLTC, PSRC1 |
| Microtubule depolymerization | 1.73 | 0.00295 | **TPX2**, MID1, FGF13, KIF2A, MID1IP1 |
| Protein transport along microtubule | 1.65 | 0.00300 | **KIF3B**, KIFAP3, RPGR, IFT22, DYNC2H1 |

### Figure. S4

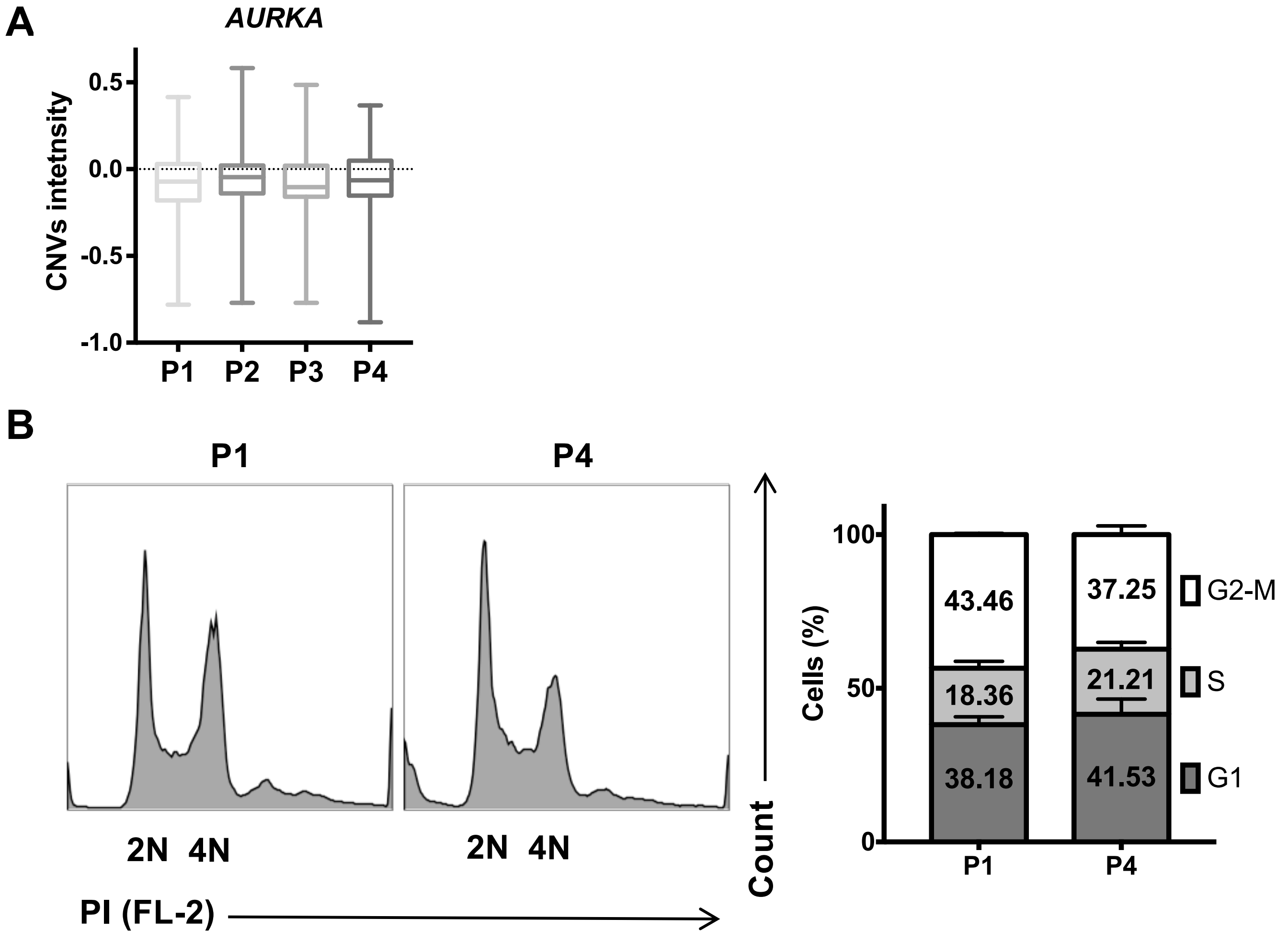
A
AURKA
CNVs intetnsity
0.5
0.0
-0.5
-1.0
P1
P2
P3
P4
B
P1
P4
2N 4N
2N 4N
PI (FL-2)
Count
Cells (%)
100
50
0
43.46
18.36
38.18
37.25
21.21
41.53
G2-M
S
G1
P1
P4
